## Supplemental material for "Neuronal network dysfunction in a human model for Kleefstra syndrome mediated by enhanced NMDAR signaling"

### Supplementary Materials

Fig. S1. Characterization of control and KS patients iPS cells.

Fig. S2. Characterization of control and KS iPS cells and iNeurons.

Fig. S3. Single cell characterization of control and KS iNeurons.

Fig. S4. Distribution of network burst duration and interval for control and KS neuronal networks.

Fig. S5. Characterization of CRIPR/cas9-edited iPS cell and iNeurons.

Fig. S6. Discriminant analysis classification of control and KS iNeurons.

Fig. S7. Effect of AMPA- and NMDA-blockage on control and KS network.

Table S1. Primers designed for RT-qPCR experiments targeting human transcripts.

Table S2. Primers designed for ChIP-qPCR experiments targeting human promoter regions.

Table S3. Overview of statistical analyses.

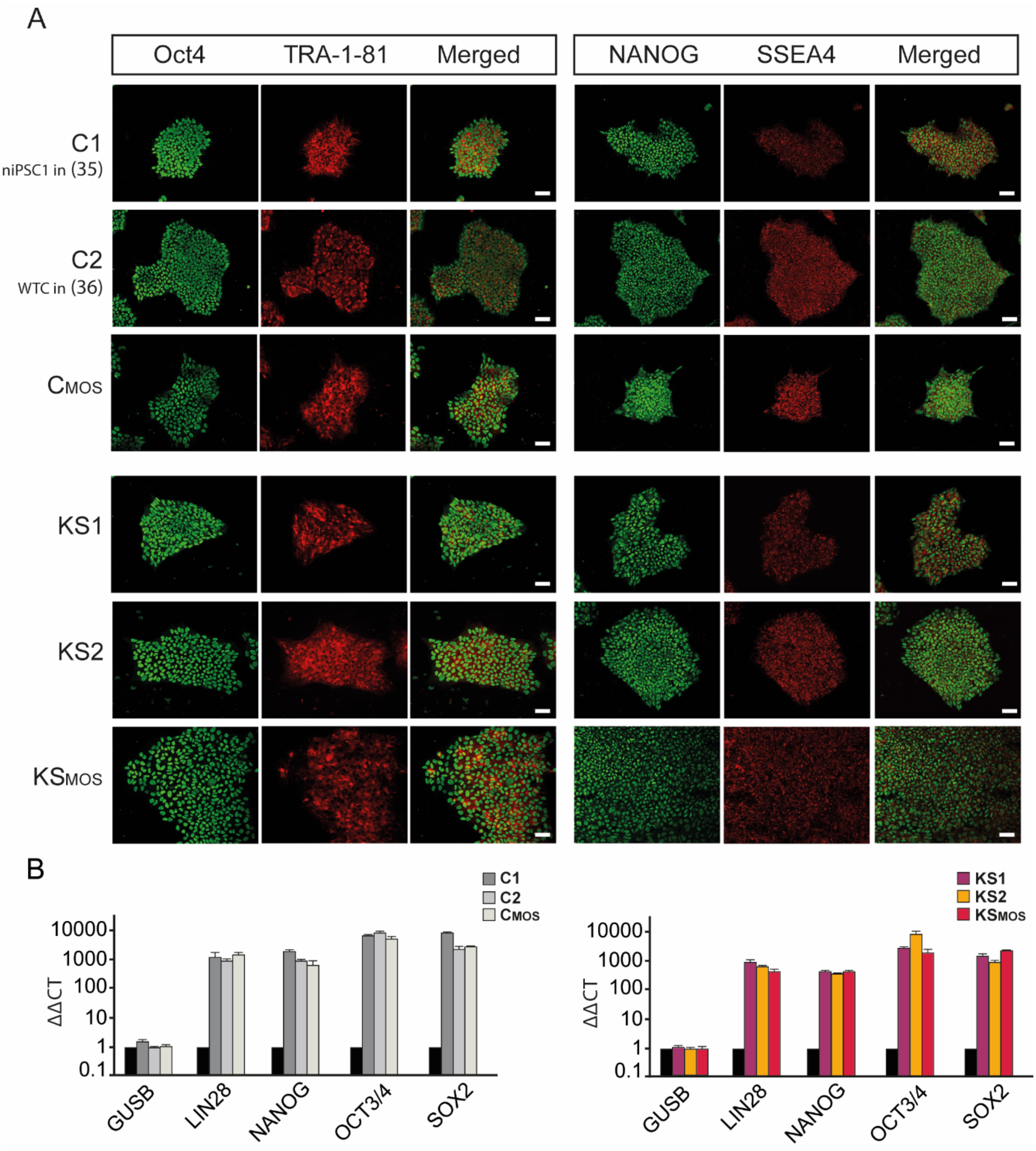


**Fig. S1. Characterization of control and KS patients iPS cells. A)** Representative images of iPS cell colonies from 3 control and 3 KS lines ($C_1$, $C_2$, $C_{MOS}$; $KS_1$, $KS_2$, $KS_{MOS}$) stained for different pluripotency marker (scale bar 50 µM). All lines used in this study were examined for the expression of the nuclear marker OCT4 and NANOG (green) and the surface marker TRA-1-81 and SSEA4 (red) by means of immunocytochemistry. **B)** QPCR data for control (left panel, $C_1$, $C_2$ and $C_{MOS}$) and KS patient (right panel, $KS_1$, $KS_2$ and $KS_{MOS}$) derived iPSCs showing an upregulation of pluripotency markers in iPS cells relative to their expression in corresponding parent fibroblasts (black).

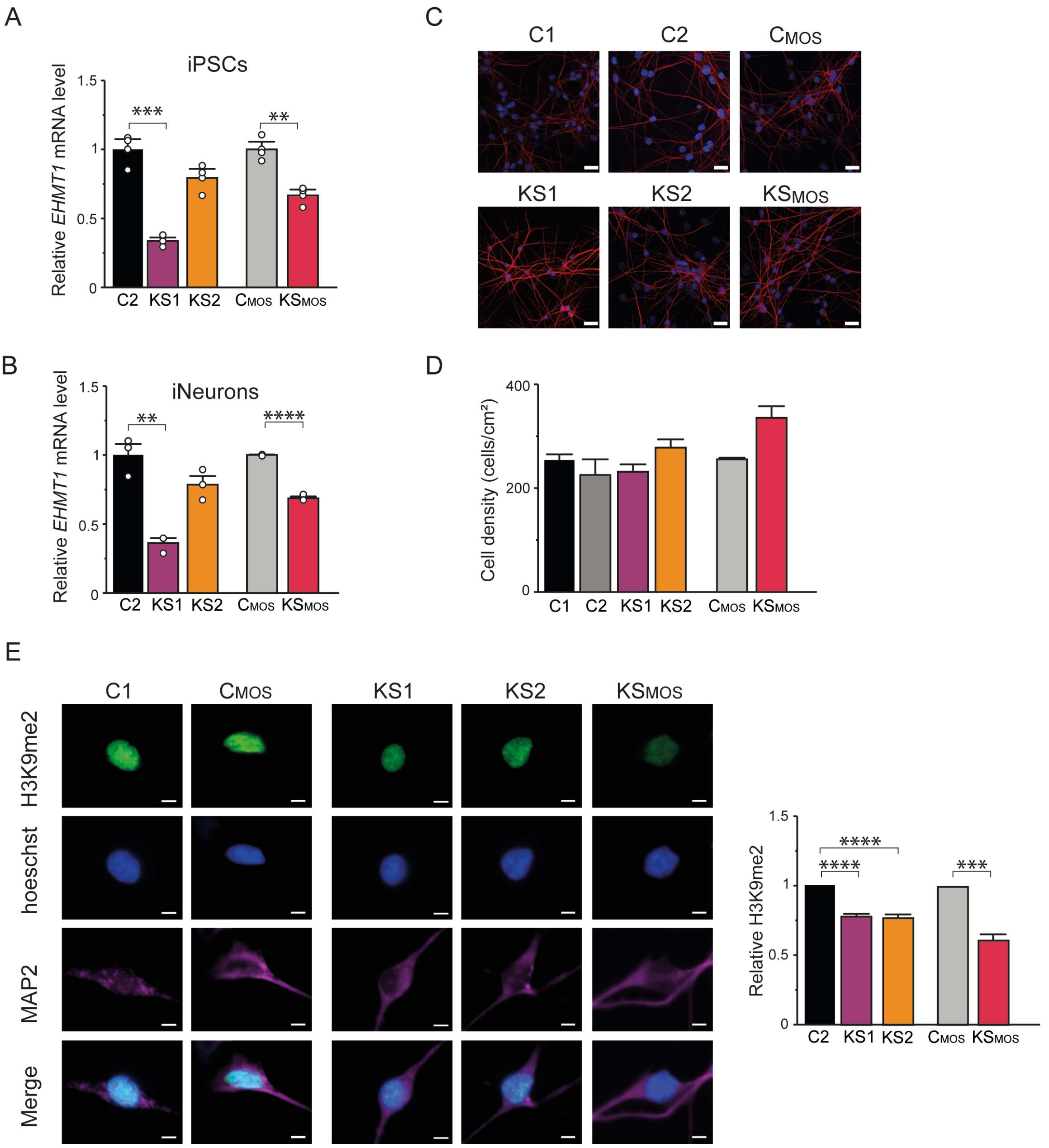


**Fig. S2. Characterization of control and KS iPS cells and iNeurons. A-B)** Graphs respectively showing relative *EHMT1* mRNA level in **A)** iPSCs and in **B)** iNeurons. Each dot represents one experiment: n=3 for all lines (i.e. controls: $C_2$, $C_{MOS}$; KS: $KS_1$, $KS_2$, $KS_{MOS}$). Values from $KS_1$ and $KS_2$ are normalized to $C_2$; values from $KS_{MOS}$ are normalized to $C_{MOS}$. **C)** Representative figure of control and KS iNeurons stained for MAP2 at DIV 21 (scale bar 50 µM). **D)** Quantification of the density of neurons derived from 3 controls and 3 KS iNeurons at 21 DIV. **E)** Representative figure of control and KS iNeurons stained for H3K9me2 (green) and MAP2 (pink) at DIV 3 (scale bar 5 µm) and quantification of H3K9me2 mean fluorescence intensity (MFI). n=38 for $C_2$; n=15 for $C_{MOS}$; n=130 for $KS_1$; n=52 for $KS_2$; n=24 for $KS_{MOS}$. Values from $KS_1$ and $KS_2$ are normalized to $C_2$; values from $KS_{MOS}$ are normalized to $C_{MOS}$. Data represent means ± SEM. ** P<0.005, *** P<0.0005, one-way ANOVA test and post hoc Bonferroni correction was performed between controls and KS-derived cultures.

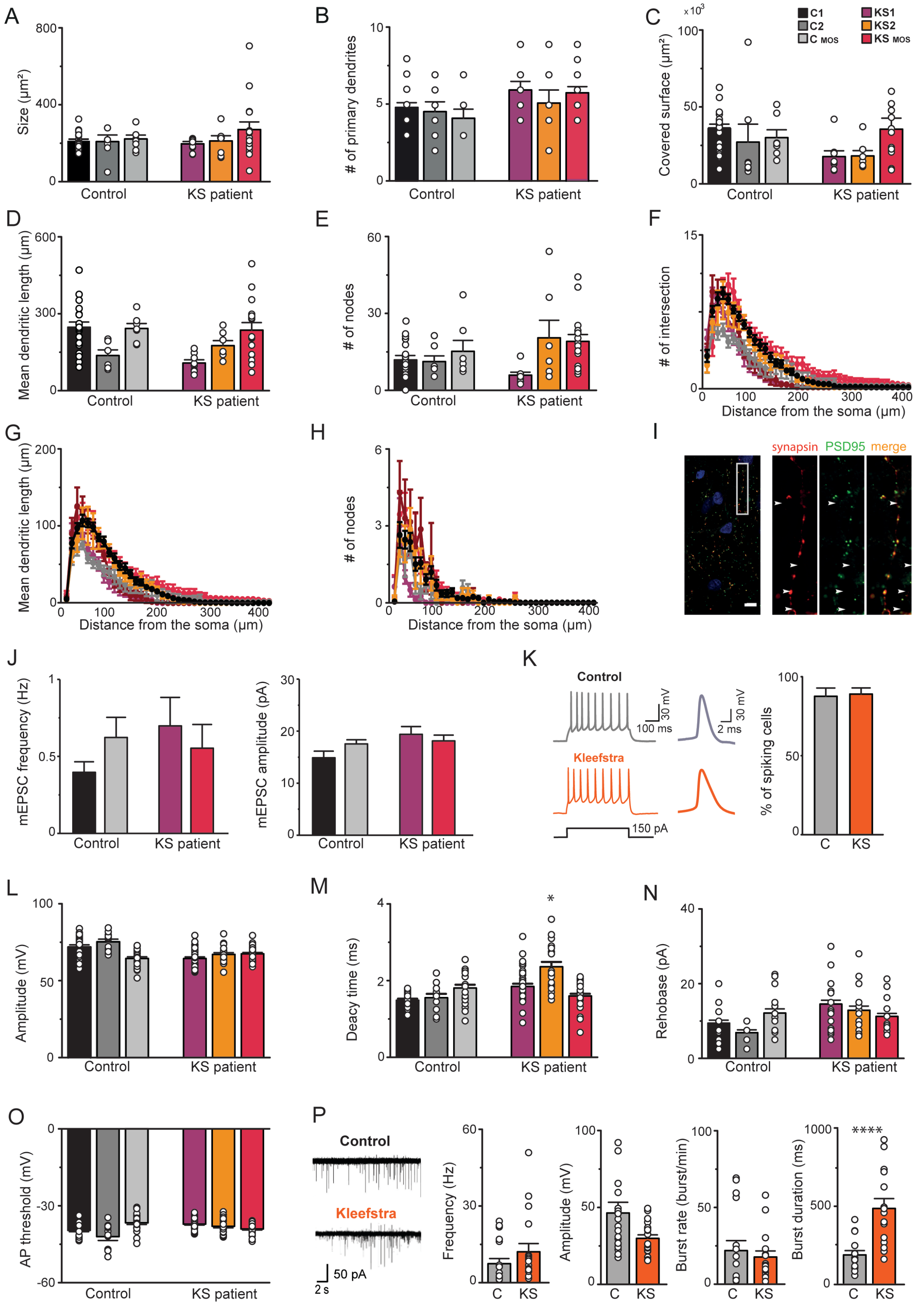


**Fig. S3. Single cell characterization of control and KS iNeurons**. **A-E)** Graphs respectively showing the **A)** neuronal size, **B)** number of primary dendrites, **C)** covered surface, **D)** mean dendritic length and **E)** number of nodes for neurons derived from 3 control and 3 KS iNeurons. **F-H)** Graph showing the Sholl analysis on **F)** number of intersections, **G)** mean dendritic length and **H)** number of nodes. Dendrite intersections were counted at 10 μm intervals. Each dot represents one cell: n=22 for $C_1$; n=7 for $C_2$; n=7 for $C_{MOS}$; n=7 for $KS_1$; n=8 for $KS_2$; n=16 for $KS_{MOS}$. **I)** Representative figure of a control iNeurons stained for synapsin 1/2 (red) and PSD95 (green) at DIV 21 (scale bar 10 μm) to indicate the formation of functional synapses with a pre- and post-synaptic site. **J)** Graph showing the frequency and amplitude of mEPSCs received by two control ($C_1$ and $C_{MOS}$) and two KS patient ($KS_1$ and $KS_{MOS}$) derived neurons. n=10 for $C_1$, n=11 for $C_{MOS}$, n=9 for $KS_1$ and n=12 for $KS_{MOS}$. **K)** Representative example traces of action potentials generated by control and KS patient iNeurons (grey and orange respectively) and graph showing the percentage of cells generating a train of 10 action potentials (n=60 for C; n=85 for KS). **L-O)** Graphs respectively showing **L)** the amplitude of the action potentials (AP), **M)** the decay time, **N)** the rheobase (i.e. current needed to generate an AP) and **O)** the AP threshold for neurons derived from 3 control and 3 KS iNeurons. Each dot represents one cell: n=24 for $C_1$; n=12 for $C_2$; n=24 for $C_{MOS}$; n=33 for $KS_1$; n=22 for $KS_2$; n=30 for $KS_{MOS}$. **P)** Example traces and quantification of whole cell voltage-clamp recordings of spontaneous excitatory postsynaptic currents (sEPSCs) in control- and KS iNeurons. Each dot represents one cell: n=16 for C; n=16 for KS. Data represent means ± SEM. * P<0.05, **** P<0.0001. One-way ANOVA test and post hoc Bonferroni correction was performed between controls and KS patient-derived cultures.

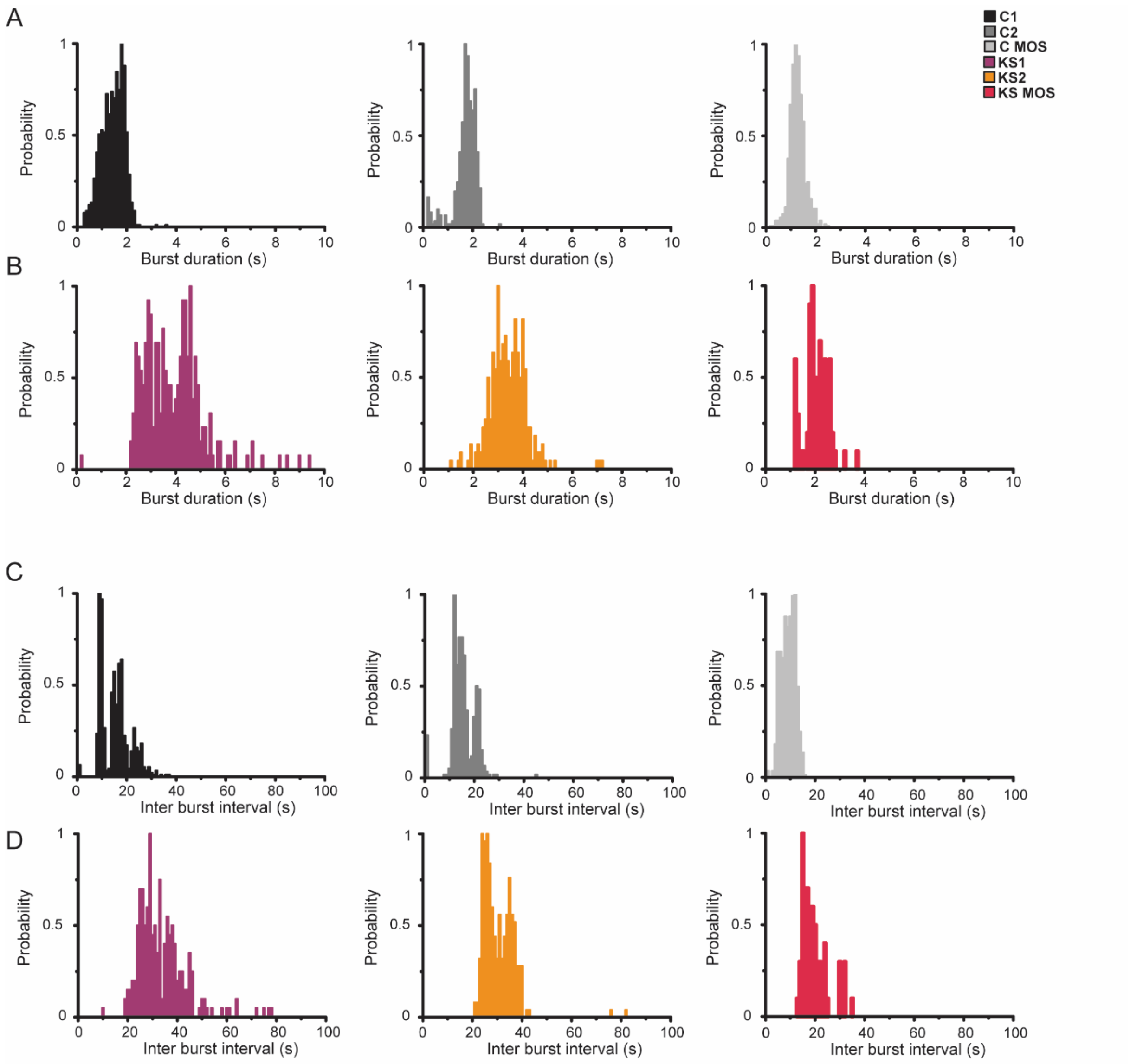


**Fig. S4. Distribution of network burst duration and interval for control and KS neuronal networks. A-B)** Graphs showing the network burst duration distribution for **A)** control- and **B)** Kleefstra patient-derived neuronal networks (bin size=100 ms) **C-D)** Graphs showing the network burst interval distribution for **C)** control- and **D)** Kleefstra patient-derived neuronal networks (bin size=1 s).

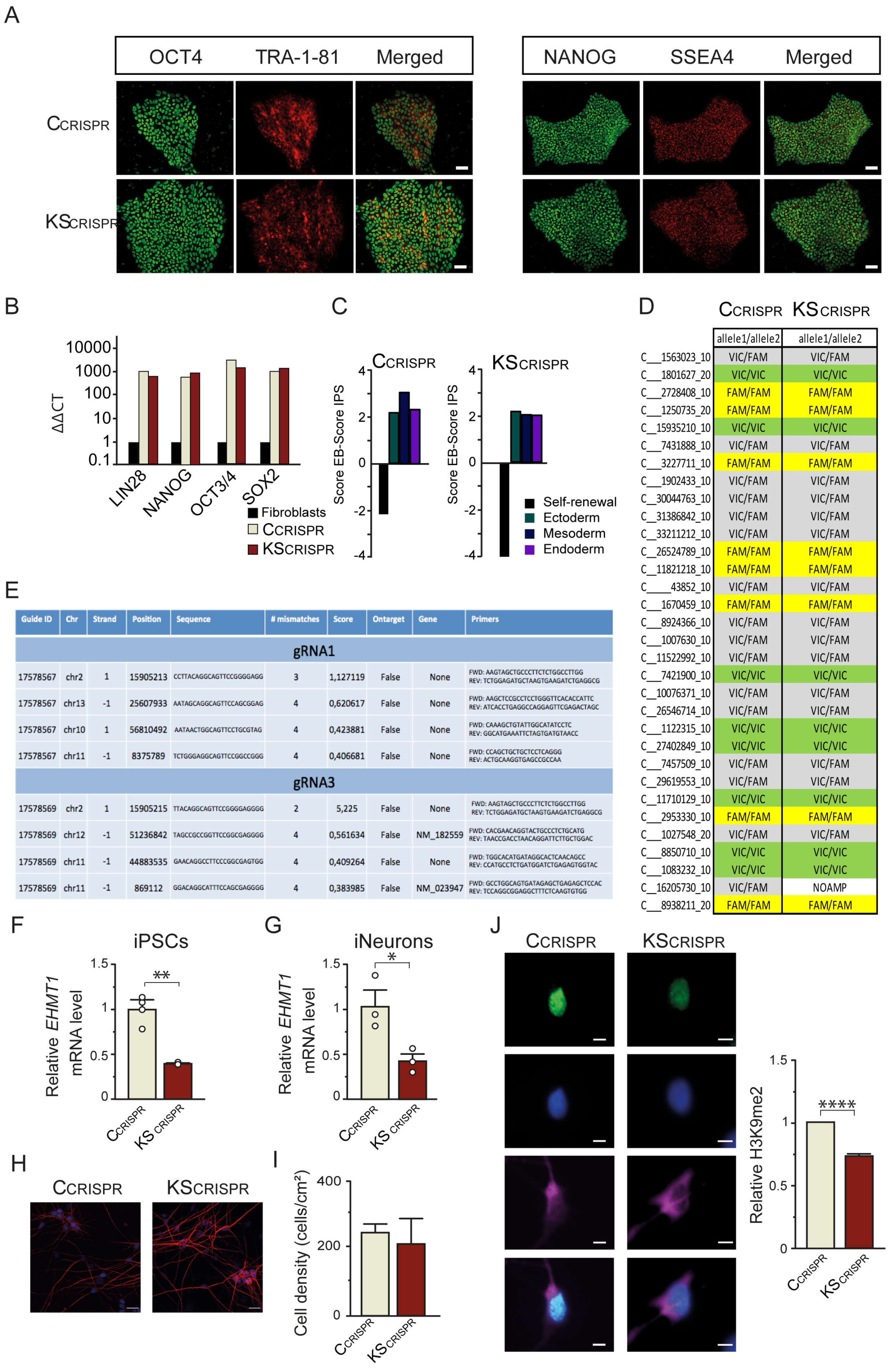


D

| | CCRISPR allele1/allele2 | KSCRISPR allele1/allele2 |
|---|---|---|
| C___1563023_10 | VIC/FAM | VIC/FAM |
| C___1801627_20 | VIC/VIC | VIC/VIC |
| C___2728408_10 | FAM/FAM | FAM/FAM |
| C___1250735_20 | FAM/FAM | FAM/FAM |
| C__15935210_10 | VIC/VIC | VIC/VIC |
| C___7431888_10 | VIC/FAM | VIC/FAM |
| C___3227711_10 | FAM/FAM | FAM/FAM |
| C___1902433_10 | VIC/FAM | VIC/FAM |
| C__30044763_10 | VIC/FAM | VIC/FAM |
| C__31386842_10 | VIC/FAM | VIC/FAM |
| C__33211212_10 | VIC/FAM | VIC/FAM |
| C__26524789_10 | FAM/FAM | FAM/FAM |
| C__11821218_10 | FAM/FAM | FAM/FAM |
| C_____43852_10 | VIC/FAM | VIC/FAM |
| C___1670459_10 | FAM/FAM | FAM/FAM |
| C___8924366_10 | VIC/FAM | VIC/FAM |
| C___1007630_10 | VIC/FAM | VIC/FAM |
| C__11522992_10 | VIC/FAM | VIC/FAM |
| C___7421900_10 | VIC/VIC | VIC/VIC |
| C__10076371_10 | VIC/FAM | VIC/FAM |
| C__26546714_10 | VIC/FAM | VIC/FAM |
| C___1122315_10 | VIC/VIC | VIC/VIC |
| C__27402849_10 | VIC/VIC | VIC/VIC |
| C___7457509_10 | VIC/FAM | VIC/FAM |
| C__29619553_10 | VIC/FAM | VIC/FAM |
| C__11710129_10 | VIC/VIC | VIC/VIC |
| C___2953330_10 | FAM/FAM | FAM/FAM |
| C___1027548_20 | VIC/FAM | VIC/FAM |
| C___8850710_10 | VIC/VIC | VIC/VIC |
| C___1083232_10 | VIC/VIC | VIC/VIC |
| C__16205730_10 | VIC/FAM | NOAMP |
| C___8938211_20 | FAM/FAM | FAM/FAM |

E

| Guide ID | Chr | Strand | Position | Sequence | # mismatches | Score | Ontarget | Gene | Primers |
|---|---|---|---|---|---|---|---|---|---|
| gRNA1 | | | | | | | | | |
| 17578567 | chr2 | 1 | 15905213 | CCTTACAGGCAGTTCCGGGGAGG | 3 | 1,127119 | False | None | FWD: AAGTAGCTGCCCTTCTCTGGCCTTGG<br>REV: TCTGGAGATGCTAAGTGAAGATCTGAGGCG |
| 17578567 | chr13 | -1 | 25607933 | AATAGCAGGCAGTTCCAGCGGAG | 4 | 0,620617 | False | None | FWD: AAGCTCCGCCTCCTGGGTTCACACCATTC<br>REV: ATCACCTGAGGCCAGGAGTTCGAGACTAGC |
| 17578567 | chr10 | 1 | 56810492 | AATAACTGGCAGTTCCTGCGTAG | 4 | 0,423881 | False | None | FWD: CAAAGCTGTATTGGCATATCCTC<br>REV: GGCATGAAATTCTAGTGATGTAACC |
| 17578567 | chr11 | -1 | 8375789 | TCTGGGAGGCAGTTCCGGCCGGG | 4 | 0,406681 | False | None | FWD: CCAGCTGCTGCTCCTCAGGG<br>REV: ACTGCAAGGTGAGCCGCCAA |
| gRNA3 | | | | | | | | | |
| 17578569 | chr2 | 1 | 15905215 | TTACAGGCAGTTCCGGGGAGGGG | 2 | 5,225 | False | None | FWD: AAGTAGCTGCCCTTCTCTGGCCTTGG<br>REV: TCTGGAGATGCTAAGTGAAGATCTGAGGCG |
| 17578569 | chr12 | -1 | 51236842 | TAGCCGCCGGTTCCGGCGAGGGG | 4 | 0,561634 | False | NM_182559 | FWD: CACGAACAGGTACTGCCCTCTGCATG<br>REV: TAACCGACCTAACAGGATTCTTGCTGGAC |
| 17578569 | chr11 | -1 | 44883535 | GAACAGGCCTTCCCGGCGAGTGG | 4 | 0,409264 | False | None | FWD: TGGCACATGATAGGCACTCAACAGCC<br>REV: CCATGCCTCTGATGGATCTGAGAGTGGTAC |
| 17578569 | chr11 | -1 | 869112 | GGACAGGCATTTCCAGCGAGGGG | 4 | 0,383985 | False | NM_023947 | FWD: GCCTGGCAGTGATAGAGCTGAGAGCTCCAC<br>REV: TCCAGGCGGAGGCTTTCTCAAGTGTGG |

**Fig. S5. Characterization of CRIPR/cas9-edited iPS cell and iNeurons. A)** Representative images of iPS cell colonies ($C_{CRISPR}$ and $KS_{CRISPR}$) stained for different pluripotency marker (scale bar 50 µM). $C_{CRISPR}$ and $KS_{CRISPR}$ iPS cells lines were examined for the expression of the nuclear marker OCT4 and NANOG (green) and the surface marker TRA-1-81 and SSEA4 (red) by means of immunocytochemistry. **B)** QPCR data for control ($C_{CRISPR}$) and CRISPR edited ($KS_{CRISPR}$) line showing an upregulation of pluripotency markers in iPSCs relative to their expression in the fibroblasts. **C)** Quantitative analysis of tri-lineage differentiation potential; both $C_{CRISPR}$ and $KS_{CRISPR}$ lines have the capacity to differentiate towards all three germ layers **D)** DNA fingerprinting: the provided genome-edited iPSC line shows identical SNP profile with the corresponding parent iPS cell line used for gene targeting. **E)** Top four off-target sites of each gRNA have been sequenced and no mutations were detected. **F-G)** Graphs respectively showing relative *EHMT1* mRNA level in **F)** iPS cells and **G)** iNeurons. Each dot represents one experiment: n=3 for $C_{CRISPR}$ and $KS_{CRISPR}$. Values from $KS_{CRISPR}$ are normalized to $C_{CRISPR}$. **H)** Representative figure of $C_{CRISPR}$- and $KS_{CRISPR}$-derived neuronal networks stained for MAP2 at DIV 21 (scale bar 50 µm). **I)** Quantification of the density $C_{CRISPR}$ and $KS_{CRISPR}$ iNeuons at DIV 21. **J)** Representative figure of control and KS iNeurons stained for H3K9me2 (green) and MAP2 (pink) at DIV 3 (scale bar 5 µm) and quantification of H3K9me2 mean fluorescence intensity (MFI) in $C_{CRISPR}$ and $KS_{CRISPR}$ iNeurons. n=36 for $C_{CRISPR}$; n=140 for $KS_{CRISPR}$. Values from $KS_{CRISPR}$ are normalized to $C_{CRISPR}$. Data represent means ± SEM. ** $P<0.005$, *** $P<0.0005$, one-way ANOVA test and post hoc Bonferroni correction was performed between controls and KS-derived cultures.

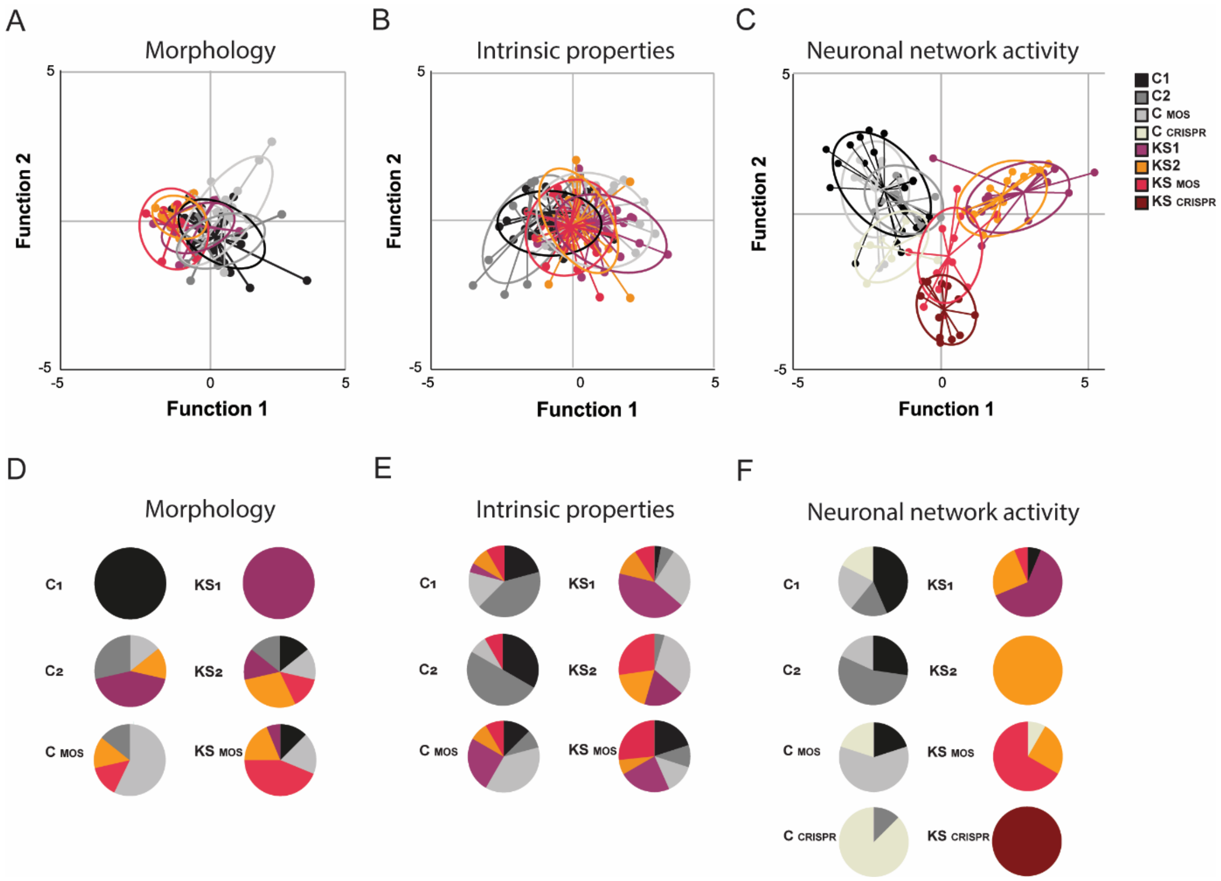


**Fig. S6. Discriminant analysis classification of control and KS iNeurons. A-C)** Discriminant function analysis with canonical discriminant functions classifies $C_1$, $C_2$, $C_{MOS}$, $KS_1$, $KS_2$ and $KS_{MOS}$ lines based on **A)** the neuronal size, the number of primary dendrites, the covered surface, the mean dendritic length and the number of nodes (i.e. morphology) (43% correct classification), **B)** the amplitude of the AP, the decay time, the rheobase and the AP threshold (i.e. intrinsic properties) (32% correct classification) and **C)** discriminant function analysis with canonical discriminant functions classifies $C_1$, $C_2$, $C_{MOS}$, $C_{CRISPR}$, $KS_1$, $KS_2$, $KS_{MOS}$ and $KS_{CRISPR}$ lines based on the firing rate, network bursting rate, network burst duration, percentage of spike outside network burst and coefficient of variability of the inter-burst interval (i.e. neuronal network activity) (70% correct classification). Group envelopes (ellipses) are centered on the group centroids. The experiment for each line and the ellipses are shown in different colors (i.e. black, dark grey, light grey and white for $C_1$, $C_2$, $C_{MOS}$, $C_{CRISPR}$ and purple, yellow, light red and dark red for $KS_1$, $KS_2$, $KS_{MOS}$, $KS_{CRISPR}$ respectively). Morphology: each dot represents one cell: n=22 for $C_1$; n=7 for $C_2$; n=7 for $C_{MOS}$; n=7 for $KS_1$; n=8 for $KS_2$; n=16 for $KS_{MOS}$. Intrinsic properties: each dot represents one cell: n=24 for $C_1$; n=12 for $C_2$; n=24 for $C_{MOS}$; n=33 for $KS_1$; n=22 for $KS_2$; n=30 for $KS_{MOS}$. Neuronal network activity: Each dot represents one experiment: n=23 for $C_1$; n=10 for $C_2$; n=10 for $C_{MOS}$; n=7 for $C_{CRISPR}$; n=15 for $KS_1$; n=15 for $KS_2$; n=12 for $KS_{MOS}$ and n=12 for $KS_{CRISPR}$. **D-F)** Pie diagrams indicating the predicted group membership for each line based on **D)** neuronal morphology, **E)** intrinsic properties and **F)** neuronal network activity.

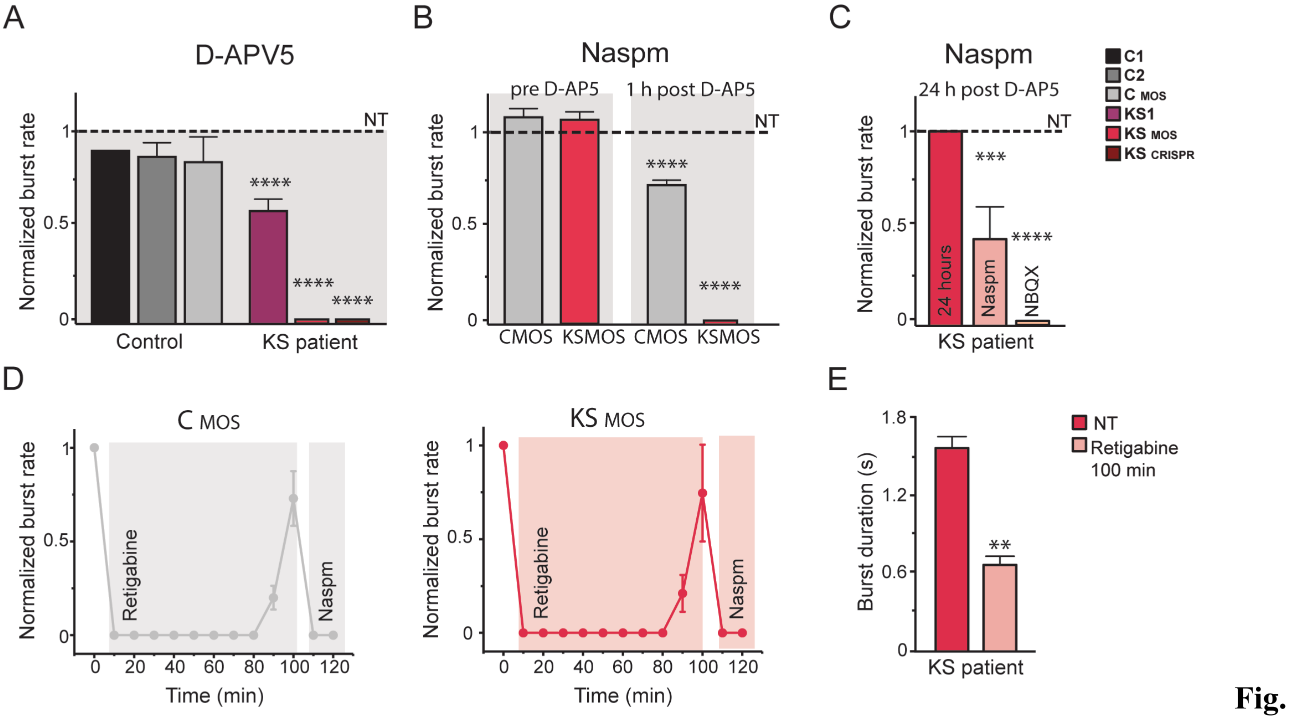


**Fig. S7. Effect of AMPA- and NMDA-blockage on control and KS network. A)** Graph showing the effect of D-AP5 (60 µM) treatment on the neuronal network burst frequency for control- and KS networks at DIV 28. The D-AP5 response (10 min after application) is shown for $C_1$, $C_2$, $C_{MOS}$, $KS_1$, $KS_{MOS}$ and $KS_{CRISPR}$. The values are normalized by the non-treated (NT) condition. The values are shown in different colors (i.e. black, dark grey, light grey for $C_1$, $C_2$, $C_{MOS}$ and purple, light red and dark red for $KS_1$, $KS_{MOS}$, $KS_{CRISPR}$ respectively). n=6 for $C_1$, n=6 for $C_2$, n=3 for $C_{MOS}$, n=10 for $KS_1$, n=14 for $KS_{MOS}$, n=14 for $KS_{CRISPR}$. **B)** Graph showing the effect of Naspm (10 µM) treatment on the neuronal network burst frequency for $C_{MOS}$ and $KS_{MOS}$ in two conditions: non-treated cultures; after 1 hour of treatment with D-AP5. The values are normalized to the non-treated (NT) condition. The values are shown in different colors (i.e. light grey for $C_{MOS}$ and light red for $KS_{MOS}$ respectively). n=6 for $C_{MOS}$ and $KS_{MOS}$ not treated; n=10 for $C_{MOS}$ and $KS_{MOS}$ D-APV treated. **C)** Graph showing the effect of Naspm (10 µM) on $KS_{MOS}$ neuronal network bursting activity 24 hours after D-AP5 treatment (NT indicates network burst activity before D-AP5 treatment). Naspm alone is not completely blocking the network bursting activity anymore. The network bursting activity is blocked when NBQX (50 µM) is added. n=3. The values are normalized to the bursting frequency 24 hours after D-AP5 treatment. **D)** Retigabine (10 µM) effect on $C_{MOS}$ and $KS_{MOS}$ neuronal network activity during time. After 100 min, Naspm (10 µM) was added in the medium and the network burst disappeared in both $C_{MOS}$ and $KS_{MOS}$. The values are normalized to the non-treated (NT) condition. **E)** Graph showing the duration of the network burst of $KS_{MOS}$ before and after 100 min of Retigabine (10 µM) treatment. n=3. Data represent means ± SEM. * $P<0.05$, *** $P<0.0005$, **** $P<0.0001$, one-way ANOVA test and post hoc Bonferroni correction was performed between controls and KS networks and Mann-Whitney test was performed between two groups.

| Human Gene Symbol | | Primer sequence 5' → 3' |
|---|---|---|
| ***EHMT1*** | Forward | GCTGGGAGAAGAGACACCTA |
| | Reverse | TGCTGGCATCGCTGTTT |
| ***GRIA1*** | Forward | GCAGCAGTGGAAGAATAGTGATG |
| | Reverse | ATCACCTTCACCCCATCGTA |
| ***GRIA2*** | Forward | GCTTGGTGCTAAATTGCTGT |
| | Reverse | TCCAAGAAAAGTAGAGCATCCA |
| ***GRIA3*** | Forward | TTCCCACTGGAGGCATGTG |
| | Reverse | CATCAGCAATATTCGTGTCATGC |
| ***GRIA4*** | Forward | TGCTGCAACTAAGACCTTCGTTAC |
| | Reverse | TCGAGTATCCCCTGTCTGTGTC |
| ***GRIN1*** | Forward | CGCCGCTAACCATAAACAAC |
| | Reverse | GGGGAATCTCCTTCTTGACC |
| ***GRIN2A*** | Forward | AGCTGCTACGGGCAGATG |
| | Reverse | CCTGGTAGCCTTCCTCAGTG |
| ***GRIN2B*** | Forward | GGAGTTCTGGTTCCTACTGGG |
| | Reverse | TCTCATGGGAACAGGAATGG |
| ***PPIA (housekeeping)*** | Forward | CATGTTTTCCTTGTTCCCTCC |
| | Reverse | CAACACTCTTAACTCAAACGAGGA |

**Table S1.** Primers designed for RT-qPCR experiments targeting human transcripts.

| **Human Promoter** | | **Primer sequence 5’ → 3’** |
|---|---|---|
| ***BDNF*_pr4** | Forward | TGT TTG GTC GGC TAG AAA GC |
| | Reverse | CCCAGATAACGTTACACCAAG |
| ***GRIN1*_pr2** | Forward | ACCCCAAGATCGTCAACATT |
| | Reverse | GGCATTGAGCTGAATCTTCC |
| ***PPIA*_pr1** | Forward | TTGGCCTGACCTAATGTTGG |
| | Reverse | TCCTATGGCTTCCCTTTCAG |

**Table S2.** Primers designed for ChIP-qPCR experiments targeting human promoter regions.
